# Microglial process dynamics enhance neuronal activity by shielding GABAergic synaptic inputs

**DOI:** 10.1101/2022.11.08.515728

**Authors:** Koichiro Haruwaka, Yanlu Ying, Yue Liang, Anthony D. Umpierre, Min-Hee Yi, Vaclav Kremen, Tingjun Chen, Tao Xie, Hailong Dong, Gregory A. Worrell, Long-Jun Wu

## Abstract

Microglia are resident immune cells of the central nervous system (CNS) and play key roles in brain homeostasis. During anesthesia, microglia increase their dynamic process surveillance and interact more closely with neurons. However, the functional significance of microglial process dynamics and neuronal interaction has remained unclear. Using *in vivo* two-photon imaging in awake mice, we discover that microglia enhance neuronal activity after the cessation of general anesthesia. Hyperactive neuron somata are directly contacted by microglial processes, which specifically co-localize with GABAergic boutons. Electron microscopy-based synaptic reconstruction after two-photon imaging reveals that microglial processes enter into the synaptic cleft to shield GABAergic inputs. Microglial ablation or loss of microglial β2-adrenergic receptors prevent post-anesthesia neuronal hyperactivity. Together, our study demonstrates a previously unappreciated function of microglial process dynamics, which allow microglia to transiently boost neuronal activity by physically shielding inhibitory inputs.

## Introduction

Microglia, resident immune cells of the central nervous system (CNS), are highly dynamic and constantly survey the brain environment (*1–3*). Microglia maintain brain homeostasis and can regulate neuronal circuits through their dynamic process surveillance and subsequent interactions with neurons. Microglia surveil neuronal activity by contacting pre- and post-synaptic elements, monitoring their activity, and pruning synapses (*4–8*). Particularly, microglia can dampen neuronal hyperactivity during seizures and epilepsy (*9, 10*). Dampening neuronal activity involves the rapid degradation of ATP into adenosine to reduce presynpatic activity (*11*). Potentially, ATP regulation most readily occurs at somatic purinergic junctions between microglial processes and neurons (*9, 12*). As complex mechanisms used by microglia to reduce neuronal hyperactivity have been elucidated, much remains unknown about the functional significance of microglia-neuron interactions during hypoactive periods, such as anesthesia. In this study, we investigated the potential for microglia to also regulate neuronal activity during periods of neuronal hypoactivity using *in vivo* two-photon imaging combined with electron microscopy. We find that microglia can transiently boost neuronal activity, which manifests in the period following anesthesia cessation. Increased neuronal activity occurs through microglial shielding of axosomatic GABAergic synapses onto excitatory neurons during the anesthesia phase.

## Results

### Neuronal hyperactivity during the emergence from general anesthesia

Seizures and epileptiform cortical activity can occur after the cessation of general anesthesia, termed the “emergence” or “recovery” period, in human patients (*13*) and in rodents (*14*). To directly test for neuronal hypersynchrony in the anesthesia emergence period, we employed *in vivo* two-photon microscopy in mice to observe neuronal Ca^2+^ activity in the awake baseline state, during the administration of isoflurane anesthesia, and during emergence from general anesthesia (**Fig. 1, A to C**). Excitatory neurons in layer II-III of somatosensory cortex were virally transfected with a genetically-encoded calcium indicator (AAV9.CaMKII.GCaMP6s). In the awake state, excitatory neurons exhibited spontaneous Ca^2+^ activity, which was largely silenced by isoflurane anesthesia (**Fig. 1, C and D; movie S1**). During anesthesia emergence, we found that most excitatory neurons (48.2%) increased their spontaneous firing above baseline levels. A small percentage of neurons reduced their firing relative to awake baseline (22.2%), and some exhibited no substantial change (29.6%; **Fig. 1, D and E**). Increased neuronal Ca^2+^ activity manifests as both greater maximum calcium signal amplitude and longer calcium transient duration, culminating in a much greater signal area, which could be observed as early as 15 minutes after the cessation of anesthesia. This period of hyperactivity gradually returned to initial, awake baseline levels over 60 minutes (**Fig. 1, F to H**). Consistent with neuronal hyperactivity during anesthesia emergence, we found that sensory perception was sensitized in this period by assessing the pain reflex threshold with von Frey filaments (**Fig. 1I**). In addition, we observed an increase in animal locomotion coinciding with the peak of neuronal hyperactivity during emergence (**Fig. 1, J and K**). These results demonstrate that anesthesia can result in a transient period of neuronal hyperactivity and behavioral sensitization during emergence.

**Figure 1:**
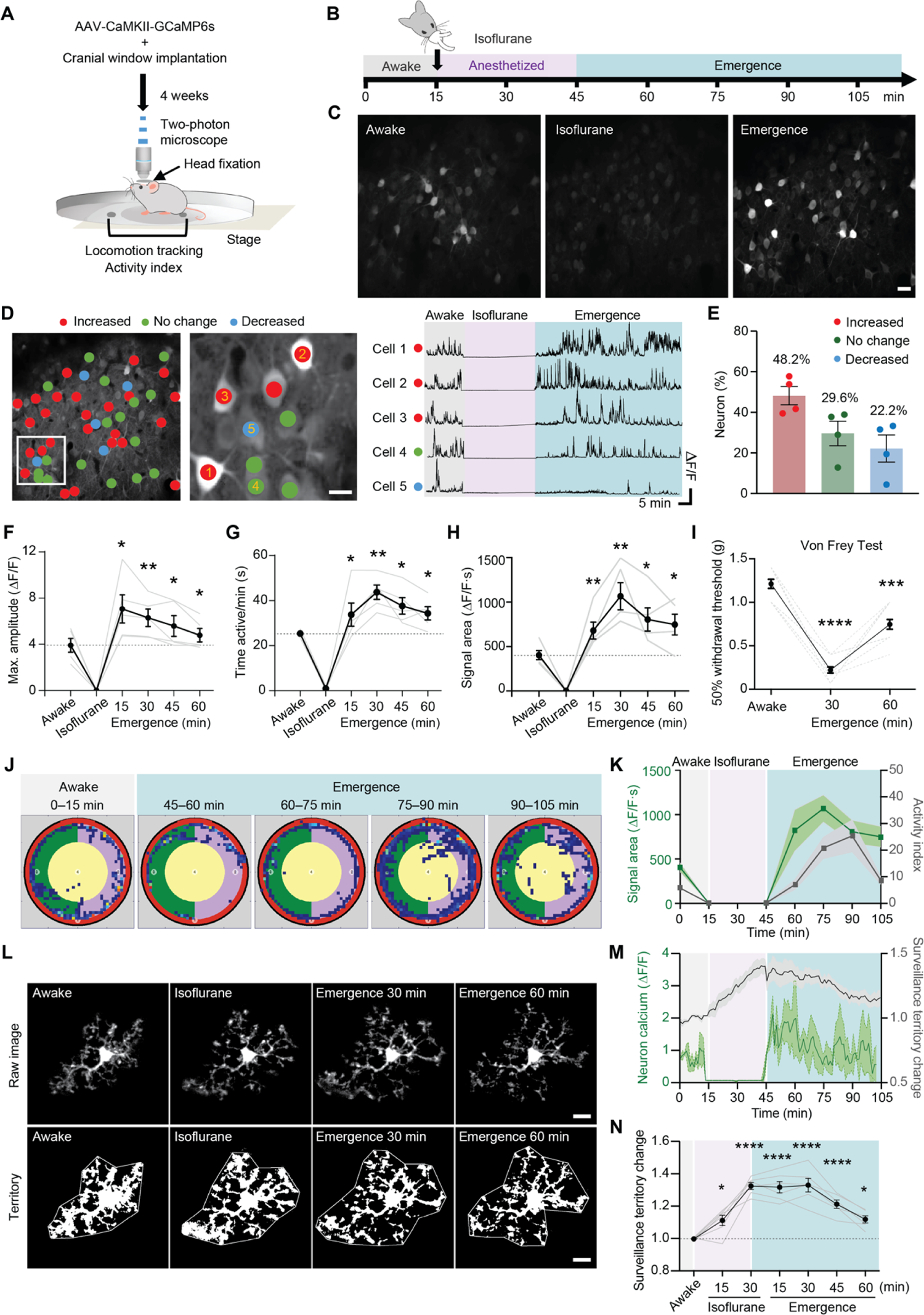
Neuronal hyperactivity occurs during emergence from general anesthesia. (**A**) Outline of *in vivo* two-photon calcium imaging with simultaneous mouse locomotion tracking. (**B**) Schematic diagram of the experimental timeline. (**C**) Representative images of neuronal calcium activity from the somatosensory cortex before, during, and after anesthesia (Scale bar: 20 μm). (**D**) Classification of neuronal calcium activity and representative ΔF/F calcium traces. Neurons with increased activity during emergence compared to awake baseline are shown in red, while those with no change are shown in green, and those with decreased activity are shown in blue. (**E**) Percentage of neurons displaying these activity changes in emergence compared to awake baseline. Exact criteria and methodology are provided in the methods (n = 4 mice). (**F-H**) Maximum amplitude (ΔF/F), active time (time of ΔF/F>0.5), and signal area (ΔF/F·s) for neuronal calcium traces at each time point. Dark line represents the mean ± SEM from n = 5 mice, while dashed lines indicate an individual animal. (**I**) Threshold for paw withdrawal in the von Frey filament test (n = 6 mice). (**J**) Representative locomotor movement of mice using a magnetic-based tracking platform. Mouse movement is displayed in blue, while all other colors distinguish platform zones. (**K**) Changes in neuronal calcium activity (mean ± SEM of ΔF/F signal area, green) and mouse locomotion activity (activity index, gray) throughout experimental phases (n = 4 mice). (**L**) Representative images of microglial morphology and surveillance territory during each experimental phase (Scale bar: 10 μm). (**M**) Time-course changes in microglial territory (gray) overlaid with corresponding changes in neuronal calcium activity (ΔF/F, green). Territory area throughout the experiment was normalized to the awake baseline start. (**N**) Time-course changes in microglial territory (n = 6 mice). In all graphs, each point or dashed line indicates data from an individual animal (**E, F, G, H, I and N**), while bars or dark lines with error bars indicate the mean ± SEM. One-way ANOVA followed by Tukey’s post-hoc test; n.s., not significant; *p < 0.05, **p < 0.01, ***p < 0.001 and ****p < 0.0001.

Microglial processes extend outward to expand their parenchymal surveillance territory during anesthesia. Anesthesia is known to reduce adrenergic tone during anesthesia by inhibiting locus coeruleus activity (*15, 16*). It is unclear how microglial processes would respond during the period of neuronal hyperactivity following anesthesia emergence. To assess microglial process dynamics across phases of anesthesia, we utilized Cx3Cr1^GFP/+^ mice. As expected, microglial territory gradually increased during isoflurane anesthesia. However, the increase of microglial territory area maintained but did not further expand during emergence (**Fig. 1, L to N; movie S2**). Thus, neuronal hyperactivity does not engender any additional increase in microglial process motility or surveillance territory during the early phase of isoflurane emergence.

### Microglia are required for neuronal hyperactivity during emergence

Greater microglia-neuron interaction under anesthesia could potentially regulate neuronal activity. The impact of these interactions could manifest as neuronal hyperactivity in the emergence period. To test this idea, we employed microglia ablation approaches combined with neuronal Ca^2+^ imaging. Mice were fed a chow containing PLX3397, a Csf1 receptor inhibitor, for 3 weeks. At the time of experimentation, approximately 98% of cortical microglia were ablated compared to wild-type mice fed a control diet (**fig. S1, A and B**). We imaged neuronal Ca^2+^ activity across experimental phases (awake baseline, anesthesia, and emergence) in mice with and without microglia (**fig. S1C**). Mice fed with the control diet still displayed neuronal hyperactivity during emergence. However, mice with ablated microglia failed to display neuronal Ca^2+^ hyperactivity during emergence (**fig. S1, D to F; movie S3**). Consistently, behavior sensitization during emergence from general anesthesia was also not observed in microglia-ablated mice (**fig. S1G**). These results suggest that microglia are required for neuronal hyperactivity during emergence from general anesthesia.

### Increased microglial contacts correlates with neuronal hyperactivity during emergence

To examine the possible correlation between microglial process dynamics and neuronal hyperactivity, we visualized neurons and their Ca^2+^ activity with GCaMP6s and tdTomato viral transfection. In addition, microglial processes were visualized using Cx3cr1^GFP/+^ mice. During anesthesia, microglial processes increased their contact with neuron somas and exhibited bulbous process endings at their contact sites with neuronal cell bodies (**Fig. 2A; movie S4**). During the awake baseline period, 83.7% of neuronal somata had one or no microglial process contact sites on their somata, and 16.3% of neuronal somata had two or more contact sites (**Fig. 2B**). During anesthesia, microglial processes contact with neuronal somas increased overall. At a population level, although some neurons retained zero or one microglial contact, similar to baseline levels (termed “low contact”), microglial processes formed two or more contacts with other neuronal somata during anesthesia (termed “high contact”). Specifically, 76.8% of neurons had high contact during anesthesia, compared to 16.3% of high contact in the awake baseline (**Fig. 2B; movie S4**). During anesthesia, the duration of microglial process contacts also increased (**fig. S2, A to C**). Bulbous endings contact with neuronal dendrites were similarly increased (**fig. S2, D to F**). In tracking neuronal activity through the various phases of imaging, we found that hyperactive neurons during emergence had the highest number of microglial contacts during anesthesia (**Fig. 2C**). In contrast, neurons with no process contact or low contact showed slower recovery from anesthesia during emergence (**Fig. 2D**). Thus, microglial contacts with neuronal somata during anesthesia strongly correlated with later hyperactivity during emergence (**Fig. 2E**).

**Figure 2:**
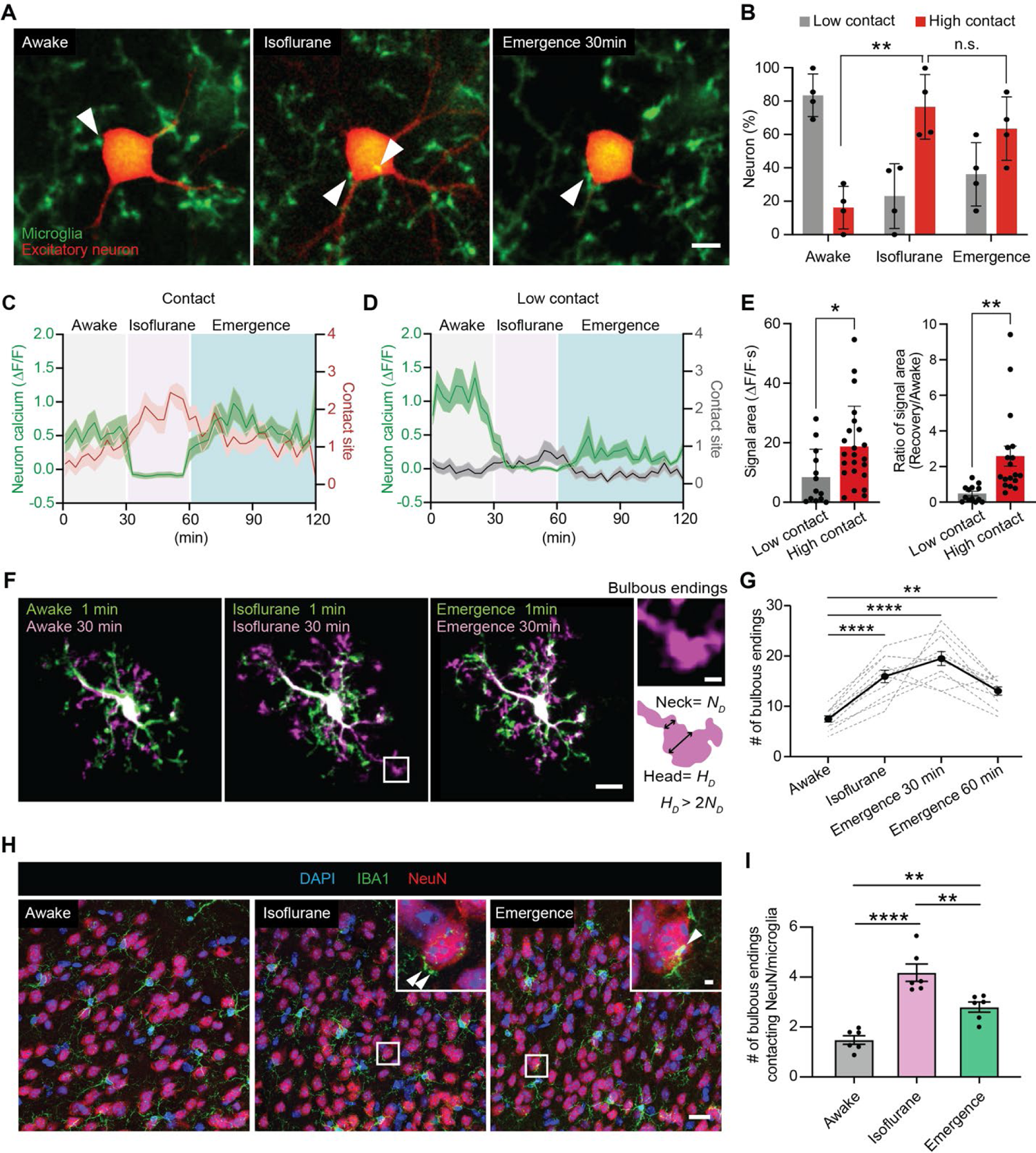
Increased microglial process contacts correlate with neuronal hyperactivity during emergence. (**A**) Representative images of microglia (green) and CaMKII neuron (red) interactions in the awake baseline, anesthesia, and anesthesia recovery periods. White arrowheads indicate contact sites (Scale bar: 5 μm). (**B**) Percentage of CaMKII neurons with soma contact from microglial processes in the awake, isoflurane, and emergence phases (n = 4 mice). (**C-D**) Overlay of GCaMP6s neuronal calcium activity (ΔF/F, green) and the number of microglial process contacts (red) with neuronal somata across anesthesia phases, separated by neurons with higher microglial contact (**C**) and lower microglial contact (**D**). (**E**) Neuronal calcium signal area in the emergence phase for somata with low or high microglial contact during anesthesia. The ratio of neuronal activity (signal area) in the awake baseline compared to anesthesia emergence. Dataset includes 13 neurons with low contact and 23 neurons with high contact from 8 mice. (**F**) Representative images of microglial processes and bulbous tip endings (white square) at the beginning (green) and end (purple) of each study phase: awake baseline, anesthesia, and emergence. Inset: Criteria for a bulbous tip ending on a microglial process (scale bar: 10 μm for main panels; 1 μm for inset). (**G**) Number of bulbous tip processes present on microglia across experimental phases. N = 11 microglia from 5 mice. (**H**) Representative immunofluorescent images of IBA1 microglia and NeuN neurons in fixed tissue prepared from mice across experimental phases. Inset shows microglial bulbous tip endings contacting NeuN somata. (**I**) Quantification of microglial bulbous tip endings contacting NeuN per microglia across experimental phases (n = 6 mice). Each point indicates data from an individual neuron (**E**) or animal (**B, I**), columns with error bars indicate the mean ± SEM. One-way ANOVA followed by Tukey’s post-hoc test (**B, G, I**), unpaired t-test (**E**); n.s., not significant; *p < 0.05, **p < 0.01, ***p < 0.001 and ****p < 0.0001.

Our results indicate that contacts by microglial bulbous endings may be important for emergence-induced neuronal hyperactivity. To further characterize the bulbous endings of microglia, we quantified these structures before, during, and after anesthesia. Bulbous endings were defined as having a process head diameter greater than twice the width of the neck portion (**Fig. 2F**). We found that microglial bulbous tip formation increased during anesthesia and peaked at 30 minutes after the cessation of anesthesia (**Fig. 2G**). We then analyzed microglial bulbous endings with neurons by Iba1 and NeuN immunostaining in layer II-III of mouse somatosensory cortex. Indeed, we found that the number of microglial bulbous endings surrounding neurons increased during anesthesia and emergence (**Fig. 2, H and I**). Together, these results demonstrate that microglia bulbous endings may represent a unique structure for neuronal interaction during anesthesia and are correlated with neuronal hyperactivity during emergence.

### Microglial processes can specifically interact with GABAergic synapses

To study how microglial contacts may promote neuronal activity, we assessed excitatory and inhibitory synapses present on a neuronal somata together with microglial processes. We used a VGAT antibody to label GABAergic synapses, VGLUT1 for glutamatergic synapses, NeuN for neuronal somata, and Iba1 for microglia in fixed cortical tissue. Interestingly, the colocalization of microglial bulbous endings with VGAT puncta was significantly increased during anesthesia and remained high in tissue fixed 30 min after anesthesia cessation (**Fig. 3, A and B; fig. S3, A to C**). Consistently, the contacts between the colocalized VGAT puncta/microglial bulbous endings with neurons were increased during anesthesia as well as emergence (**Fig. 3C**). We further quantified microglial process interactions with excitatory VGLUT1 puncta and found they were unaltered under anesthesia compared to baseline (**fig. S3, D and E**). In addition to the contacts between microglial processes and neuronal somata, we also examined soma-soma interactions between microglia and CaMKII neurons, which are also known to regulate neuronal activity during developmental seizures (*17*). However, we did not observe any change in soma-soma interactions before or after anesthesia (**fig. S3, F and G**). We then assessed whether microglia processes may phagocytose inhibitory inputs. To this end, we counted VGAT puncta but found no change in the number of VGAT puncta 24 hours after anesthesia compared to baseline, suggesting that GABAergic synapses are not removed or pruned by microglia (**Fig. 3D**). Together, these results suggest that microglia may promote neuronal activity by interacting with GABAergic inputs without synapse elimination.

**Figure 3:**
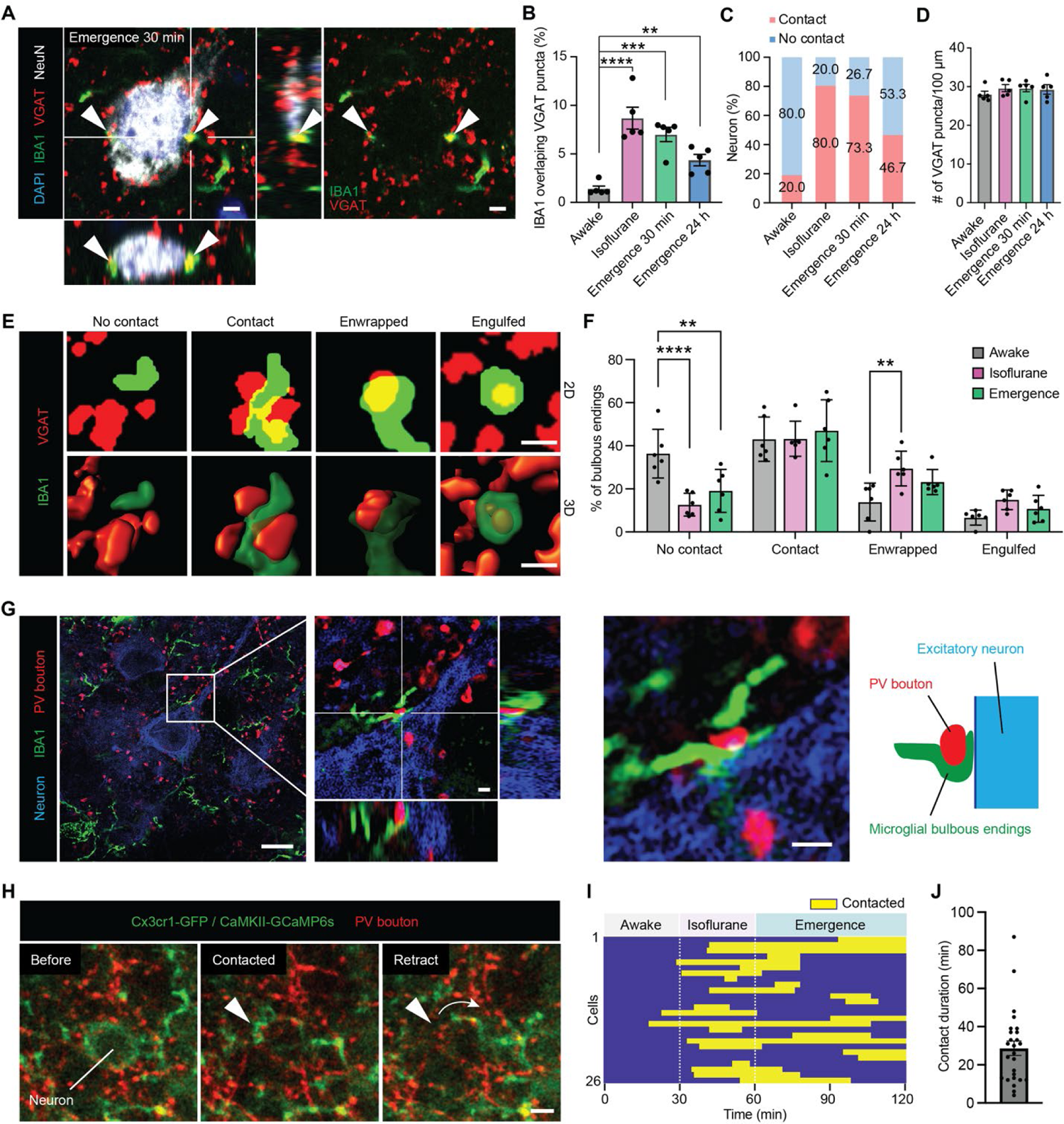
Microglial processes interact with GABAergic boutons. (**A**) Representative immunofluorescent image of DAPI nuclei (blue), IBA1 microglial processes (green), NeuN nuclei (white), and VGAT puncta (red) in cortical tissue prepared 30 min after anesthetic emergence. White arrows in main panel and orthogonal panels indicate sites of microglial bulbous tip contact with VGAT terminals (Scale bar: 2 μm). (**B**) Percentage of neurons with contact or no contact by microglial processes across experimental phases. (**C**) Percentage of IBA1 processes co-localizing with VGAT puncta across experimental phases (n = 5 mice). (**D**) Number of VGAT puncta per unit area across experimental phases (n = 5 mice). (**E**) Representative thresholded images or 3D space-filling models of microglia-VGAT interactions, including “No contact,” “partial contact,” “enwrapping,” or “full engulfment” (Scale bar: 2 μm). (**F**) Percentage of microglial bulbous tip endings engaging in different types of PV interactions across experimental phases (n = 5 mice). (**G**) Representative super-resolution image of neurons (blue), IBA1 microglia (green), and PV boutons (red) (Scale bar: 10 μm in main panels, 1 μm in inset). Examples of microglial bulbous tip interactions with PV boutons across experimental phases. Kymograph displays microglial process contact time with CaMKII neuronal somata across experimental phases. (**J**) Distribution of contact time across the whole experiment (n = 26 neurons from 5 mice). Dots represent individual mice (**C, D, F**) or individual neurons (**J**). Columns and error bars represent the mean ± SEM. One-way (**C, D**), or two-way (**F**) ANOVA followed by Tukey’s post-hoc text. *p < 0.05, **p < 0.01, ***p < 0.001 and ****p < 0.0001.

To further investigate microglial interactions with GABAergic synapses, we took a closer look at how microglial bulbous endings interact with VGAT puncta during anesthesia using confocal microscopy with high magnification Z-stacks. We classified interactions into four subtypes: VGAT puncta with no microglial contact (No contact), puncta with under 50% surface contact (Low Contact), contact with more than 50% surface area coverage (Enwrapped), and complete internalization (Engulfed) (**Fig. 3E**). This analysis indicates that the percentage of fully engulfed VGAT puncta was rare; however, the percentage of VGAT puncta enwrapped by microglia increases during anesthesia (survey of n = 420 bulbous endings from 5 mice) (**Fig. 3F**). Next, we used a combination of super-resolution microscopy and two-photon live imaging to visualize the direct interactions between microglia and inhibitory synapses. To this end, we transfected AAV9.CaMKII.GCaMP6s and AAV9.Syn.FLEX.tdTomato into parvalbumin (PV)^Cre/+^: Cx3cr1^GFP/+^ mice to simultaneously visualize neuronal Ca^2+^ activity, PV boutons, and microglia. Super-resolution microscopy confirmed that the tip of microglial bulbous endings wrapped around a portion of the PV positive bouton rather than engage in complete engulfment (**Fig. 3G**). Interestingly, two photon live imaging further revealed that the interaction of microglial bulbous endings with PV boutons was temporary and reversible (**Fig. 3H; movie S5**). During the recovery phase, bulbous endings were retracted, and PV boutons remained intact. These results indicate that microglial processes engage in the partial enwrapping of GABAergic inputs during anesthesia.

### Electron microscopy reconstruction of live imaging regions reveals shielding of PV boutons

Based on the increased microglial process contact with GABAergic boutons, we hypothesized that microglial bulbous endings may promote neuronal activity via temporarily block inhibitory inputs. Consistent with this hypothesis, we found that contacts between microglial bulbous endings and PV boutons are strongly associated with neuronal hyperactivity during the emergence from anesthesia under two photon *in vivo* imaging (**Fig. 4, A and B**). Conversely, neurons that did not have microglial contact with PV boutons at their somata during anesthesia showed little hyperactivity during post-anesthetic recovery. Thus, the percentage of hyperactive neurons during emergence is significantly increase in those with microglial contact with PV boutons than those without the contact (**Fig. 4B**).

**Figure 4:**
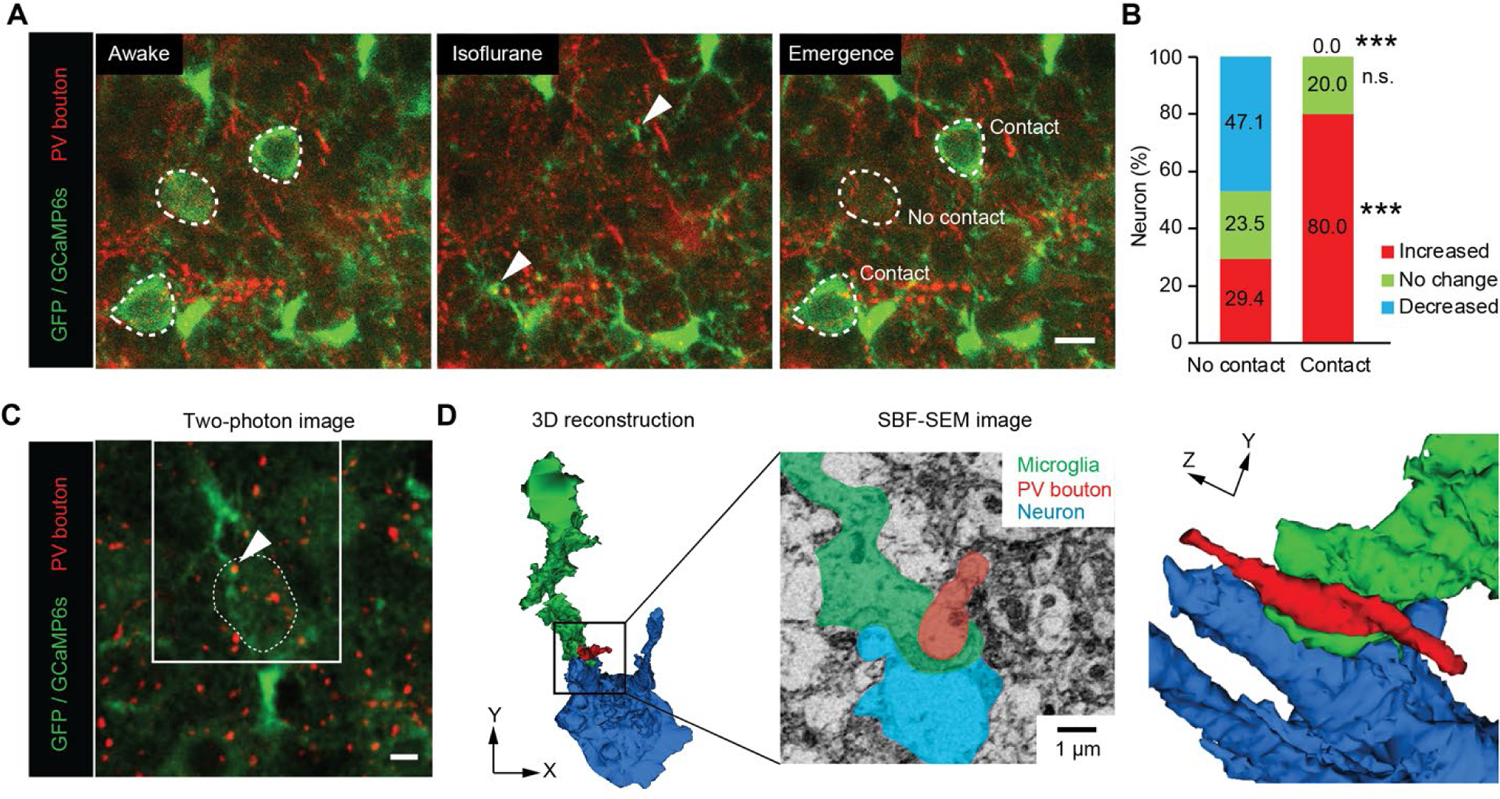
*In vivo* two-photon imaging followed by electron microscopy reconstruction reveals shielding of PV boutons. (**A**) Images of tdTomato-labelled PV boutons (AAV9.FLEX.tdTomato with PV^Cre^), microglia (Cx3cr1^GFP/+^), and neurons (AAV-CaMKII-GCaMP6s) from *in vivo* two-photon microscopy during awake, anesthesia, and emergence (Scale bar: 10 μm). (**B**) Percentage of neuronal somata displaying activity changes in emergence versus baseline as a function of microglial contact with PV boutons (n = 27 neurons from 7 mice. Chi-squared test: n.s., not significant; ***p < 0.001) (**C**) Two-photon *in vivo* imaging of microglia-PV bouton interactions adjacent to a neuronal soma, which was followed by electron microscopy reconstruction. (**D**) 3-D serial reconstruction of a microglial process, PV bouton, and neuron at the ultrastructural level using SEM.

To investigate microglial bulbous endings and their contact with PV boutons at the ultrastructural level, we performed two-photon live imaging followed by electron microscopy. Clear examples of microglia-PV interactions under *in vivo* two-photon imaging were marked via laser branding (*18*) (**Fig. 4C, fig. S4**). The ultrastructure of these interactions was investigated at the nanoscale level using serial block-face scanning electron microscopy (SBF-SEM). 3-dimensional serial reconstruction indicates that the process tips of microglia can insert themselves into a space between neuronal somata and PV boutons (**Fig. 4D; movie S6**). Thus, microglial bulbous endings were placed in a position to physically shield GABAergic synaptic inputs onto target neurons, promoting neuronal excitability by disengaging axo-somatic inhibitory inputs.

### Microglial β2 adrenergic receptors control neuronal hyperactivity during emergence

To clarify the mechanism underlying microglia’s ability to interact with axosomatic PV inputs, we investigated norepinephrine (NE) signaling. NE signaling to microglial β2-adrenergic receptor is known to restrain microglial process dynamics during wakefulness. Under anesthesia, NE tone is likely reduced, allowing microglial processes to enhance their motility and territory surveillance (*15, 16*). We first directly examined changes in NE tone during anesthesia and emergence in real time using an NE biosensor (AAV.hSyn.GRAB.NE2h; **Fig. 5A**)(*19*). Indeed, NE concentrations decreased in the parenchyma of Layer II/III somatosensory cortex during anesthesia. During the emergence from anesthesia, NE levels undergo a strong rebound in the first 30 minutes after isoflurane cessation (**Fig. 5, B and C; movie S7**). The β2-adrenergic receptor (Adrb2) is highly expressed by microglia and is a key mediator of microglial dynamics during anesthesia (*15, 16*). Accordingly, increased microglial process dynamics are lost during anesthesia in microglia-specific Adrb2 conditional knockout mice (cKO). This loss of increase in process dynamics was seen during both anesthesia and emergence (**Fig. 5, D to G; movie S8**). In addition, the directness of the process movement was increased by anesthesia in WT mice, whereas the directness was already high during the awake phase and was not altered by anesthesia in Adrb2 cKO mice (**fig. S5, A and B**). Consistently, we also observed that hypersensitivity in the von Frey test during emergence was also abolished in microglial Adrb2 cKO mice (**Fig. 5H**).

**Figure 5:**
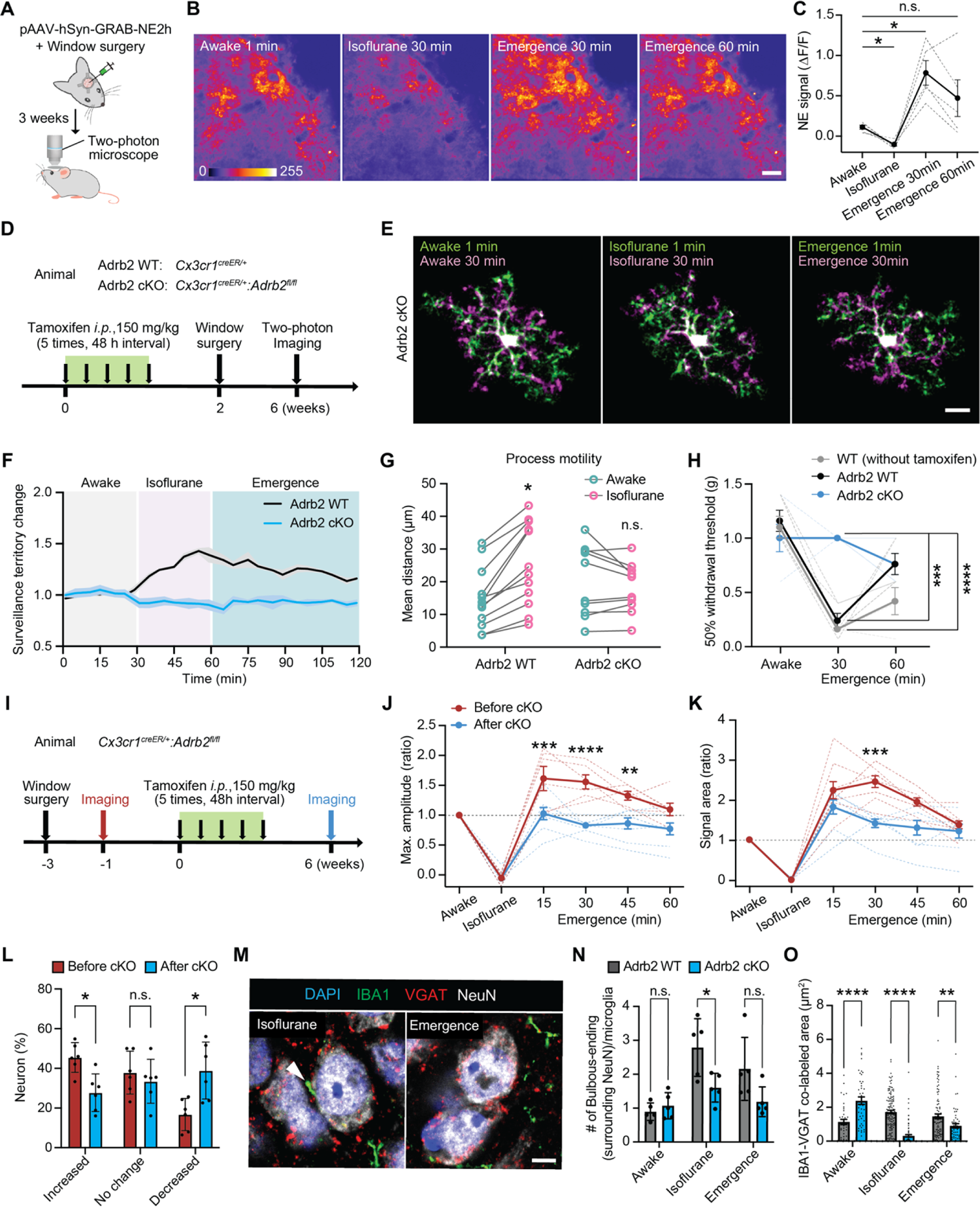
Microglial β2-adrenergic receptors in neuronal hyperactivity during emergence. (**A**) Schematic of local NE sensor transfection and cranial window surgery in the somatosensory cortex. (**B**) Representative two-photon imaging of NE biosensor fluorescence in cortex during the awake baseline, anesthesia, and emergence (Scale bar: 20 μm). (**C**) Quantification of NE sensor fluorescence (ΔF/F) across time periods (n = 5 mice, One-way ANOVA followed by Tukey’s post-hoc test). (**D**) Experimental design for comparing microglial Adrbs cKO mice (Cx3cr1^CreER/+^; Adrb2^fl/fl^) to controls (Cx3cr1^CreER/+^; Adrb2^wt/wt^) after tamoxifen recombination. (**E**) Representative images of Adrb2 cKO microglial processes at the beginning (green) and end (purple) of each study phase: awake baseline, anesthesia, and emergence (Scale bar: 10 μm). (**F**) Microglial process surveillance territory changes from awake baseline in Adrb2 cKO and WT control animals across experimental phases. (**G**) Average microglial process outgrowth/extension during the awake and anesthesia periods for Adrb2 cKO and WT controls (n = 10 WT microglia and n = 12 cKO microglia from 4 mice). (**H**) 50% paw withdrawal thresholds for C57 WT mice, Adrb2 cKO mice, and Adrb2 WT control mice during von Frey filament testing in the awake baseline and anesthesia emergence periods (n = 4 C57BL/6 WT, n = 5 Adrb2 WT, n=5 Adrb2 cKO mice). (**I**) Within-subject study design comparing responses in Cx3cr1^CreER/+^; Adrb2^fl/fl^ mice before and after tamoxifen-induced recombination. (**J-K**) Neuronal calcium signal amplitude (**J**) and signal area (**K**) across experimental time points for mice before and after recombination for Adrb2 cKO (n = 6 mice). (**L**) Percentage of neurons showing an increase, decrease, or no change in activity during emergence period compared to awake baseline. In this case, the same populations of neurons are compared before mice undergo cKO and again after tamoxifen-induced recombination (n = 6 mice). (**M**) Immunostaining of microglial processes, NeuN somata, and VGAT puncta in cKO mice (Scale bar: 5 μm). (**N**) Quantification of microglial bulbous tip endings contacting NeuN across experimental phases by genotypes (n = 5 mice). (**O**) Area of co-labelling between IBA1 microglia and VGAT puncta across experimental phases and between genotypes (n = 5 mice). In all graphs, each point or dashed line indicates data from an individual animal (**C**, **G**, **H**, **J**, **K**, **L** and **N**), or a microglial process (**O**), while columns or solid lines with error bars indicate the mean ± SEM. Two-way ANOVA followed by Tukey’s post-hoc test; n.s., not significant; *p < 0.05, **p < 0.01, ***p < 0.001 and ****p < 0.0001.

We further determined if microglial β2-adrenergic signaling underlies the regulation of neuronal hyperactivity during anesthesia and emergence. Specifically, we performed imaging experiment in two phases (**Fig. 5I**). We first imaged microglia-neuron dynamics before tamoxifen-induced Adrb2 deletion in microglia. We then performed a second round of imaging after tamoxifen-induced cKO to establish within subject controls (**Fig. 5I**). Before microglial Adrb2 deletion, we consistently observed neuronal hyperactivity during emergence, similar to levels observed in other strains. In studying the same group of neurons after Adrb2 cKO, the previous emergence-induced hyperactivity was abolished (**Fig. 5, J and K; movie S9**). At the population level, the percentage of neurons exhibiting hyperactivity during emergence decreased with microglial Adrb2 cKO (before Adrb2 cKO: 45.5%, after Adrb2 cKO: 27.7%; Fig. 5L). At a larger scale, EEG recording across experimental phases indicate that alpha, beta, and theta band power increased during emergence before Adrb2 deletion but this shift in EEG power was later lost after Adrb2 deletion (**fig. S6, A and B**). Furthermore, the number of bulbous endings around neuronal somata and their co-localization with VGAT puncta were both significantly reduced in Adrb2 cKO mice compared to WT mice during anesthesia (**Fig. 5, M to O**). Together, these results suggest that NE signaling to microglial β2-adrenergic receptors is critical to promote neuronal hyperactivity and hypersensitivity during emergence.

## Discussion

Microglia are highly dynamic immune cells critical for brain homeostasis. Microglia dampen neuronal hyperactivity during seizures and epilepsy (*9, 11*). However, it is completely unknown whether microglia influence neuronal activity under conditions like general anesthesia. Here we report that dynamic microglial processes interact with neurons and shield axo-somatic inhibitory inputs to transiently promote neuronal hyperactivity during emergence from anesthesia. Our study provides intriguing evidence that microglia can boost neuronal activity during anesthesia. Our study is part of a larger picture which indicates microglia can both negatively and positively regulate neuronal activity in a way that is homeostatic: boosting neuronal activity following hypoactivity, or dampening neuronal activity following hyperactivity.

### Microglia as a key player in synaptic remodeling

Typical roles for microglia-synapse interactions include synaptic pruning and promoting spine formation, resulting in relatively persistent changes to neuronal circuits. Synapse pruning by microglia was observed in brain development or disease contexts (*4, 5, 20*). Spine formation by microglia involves either physical contact of microglial processes with dendrites or release of BDNF from microglia (*21, 22*). In the current study, we reveal a different type of microglia-synapse interaction which regulates neuronal activity by shielding synaptic inputs without phagocytosis. A transient interaction between microglia and neurons during 30 minutes of anesthesia allows microglia to selectively displace inhibitory synapses from excitatory neuron somata. Specifically, super-resolution microscopy reveals that microglial processes selectively shield GABAergic inputs from the soma. These observations are confirmed in 3-dimentional electron microscopy (“serial reconstructions”) prepared from a region visualized *in vivo* using two-photon microscopy. This more localized axo-somatic interaction site of interaction suggests a mode of regulation different from synaptic stripping or microglia-neuron soma-soma interactions (*17, 23*). Together, our current study uncovers a novel interaction between microglial process and inhibitory boutons to transiently boost excitatory activity.

### How microglia promote neuronal activity

Recently, it has been reported that microglia reduce neuronal activity by degrading ATP and converting it into adenosine (*11*). Microglial processes also form purinergic-somatic connections with neurons in a P2Y12 receptor-dependent manner, which can also dampen neuronal activity (*9, 12*). Such studies propose microglia as a ‘brake’ on neuronal hyperactivity, playing a homeostatic or neuroprotective role in this context. In contrast, our study finds that microglia are able to enhance neuronal activity during the emergence from general anesthesia. Mechanistically, we find that microglia-neuron interactions during anesthesia promote later neuronal hyperactivity in emergence by transiently shielding inhibitory GABAergic inputs on neuronal cell bodies. Our study strongly supports the hypothesis of transient PV bouton displacement by microglia and consequent blockade of inhibitory inputs, rather than phagocytosis.

Confocal imaging reveals that an average of 30 VGAT-positive inputs synapse onto Layer II/III neuronal somata in a single plane of imaging. Microglial bulbous endings overlapped with 1.4% of these GABAergic inputs during the awake state and 8.7% of boutons during anesthesia. While interactions with GABAergic inputs may seem infrequent, axosomatic GABAergic synapses are known to exert an outsized role in controlling excitatory neuronal activity (*24*), such that relatively infrequent shielding could still have a marked physiological impact. The question remains as to how microglia selectively increase their interactions with inhibitory synapses but not excitatory synapses during anesthesia. Microglia may be able to sense GABAergic signaling directly in this period. Microglial GABA_B_ receptors have been reported to mediate inhibitory synapse pruning during development (*25*). Existing RNAseq databases also suggest these same receptors are present on microglia in adulthood (*26*). Microglia can also sense astrocyte-mediated synaptic GABA release through their GABA_B_ receptors in development (*27*). The interaction between GABAergic synapses and microglia as informed by GABA - GABA_B_ receptors on microglia in the adult brain is a key future direction.

NE has been reported as a signal to restrict microglial process motility and extension during wakefulness (*15, 16*). Under anesthesia, NE reduction is thought to allow microglial process elongation through disinhibition, thereby increasing the opportunity for physical contact between microglia and neurons. In Adrb2 cKO microglia, microglial processes were already elongated in the awake baseline period, due to a lack of NE signal reception. Adrb2 cKO prevented any changes in microglial motility or territory area during anesthesia or in the post-anesthesia recovery period. Interestingly, co-localization of microglial processes with VGAT puncta was higher in Adrb2 cKO mice than WT mice at baseline. However, in the baseline period we did not find that this influenced neuronal calcium activity or EEG power between genotypes. There was also no difference in sensory thresholds between control and cKO mice in the von Frey tests at baseline, suggesting that multiple key aspects of physiology studied herein are not radically altered by baseline differences in the microglial Adrb2 cKO animals.

### The functional significance of microglia in promoting neuronal activity

During the post-anesthesia recovery period, hypersensitivity and cognitive dysfunction are known to occur transiently in patients and may present as temporary delirium. The primary mechanisms thought to underlie this phenomenon are neurotransmitter imbalance, inflammation, and physiological stress (*13, 28*). Here we discovered an unappreciated function of microglia in promoting neuronal hyperactivity during emergence from general anesthesia. Previous *in vivo* experiments in rodents and clinical observations in patients have suggested that prolonged anesthesia can increase levels of inflammatory cytokines in the brain, with associated memory deficits (*28, 29*). In the present study, EEG power spectral analysis indicates that mice experience increased theta and delta waves during anesthesia emergence, which is often correlated with the presentation of delirium in patients (*30*). Interestingly, EEG analysis using microglial Adrb2 cKO mice indicates that alpha, beta, and theta band power do not increase during anesthesia emergence, indicating a circuit-level effect of microglia on neuronal activity. This observation co-occurs with a lack of hypersensitivity in cKO mice. The loss of microglia-inhibitory synapse interactions in Adrb2 cKO mice during anesthesia prevents PV synapses from being shielded, likely allowing GABAergic interneurons to better control/suppress hyperactivity in emergence. From our results, it would be intriguing to know if the use of β-blockers (either administered in some anesthetic regimens or taken by patients) correlates with less delirium or reduced hypersensitivity upon anesthesia emergence. Systemic administration of β-blockers can directly inhibit the microglial β2-adrenergic receptor, as we have previously demonstrated in mice (*15*).

During sleep, NE concentration in the parenchyma is reduced (*31*), similar to our observations under anesthesia. Previous studies suggest that microglia prune excitatory synapses during sleep, and chronic sleep deprivation can result in microglial reactivity/activation (*32, 33*). We found that microglia depletion (PLX3397) suppressed neuronal hyperactivity in emergence and reduced hypersensitivity. Microglia depletion can also alter sleep patterns by promoting non-rapid eye movement (NREM) sleep and decreasing nocturnal arousal (*34*). Decreased NE release during sleep might lead to microglial process extension and increased surveillance territories (*15, 16, 35*). Although there are fundamental differences between the mechanisms of anesthesia and sleep (*36*), these NE-dependent microglial functions are potentially important in a number of physiological states such as the sleep cycle. Thus, studying the broader applicability of our findings in anesthesia warrant future exploration, particularly in assessing sleep cycles.

In summary, we found that microglia transiently enhance neuronal activity to potentially counteract strong neuronal hypoactivity during general anesthesia. A novel interaction between microglial processes and GABAergic synapses shields inhibitory inputs, thereby promoting hyperexcitability. Our study demonstrates a previously unappreciated function of microglia in a positive feedback response to neuronal hypoactivity, which manifests at the cell, systems, and behavioral levels during the emergence from general anesthesia.

## Acknowledgments

This work is supported by the National Institutes of Health (R01NS088627 and R01NS112144 to L.-J.W.). Dr. Yanlu Ying is supported by National Natural Science Foundation of China (Grant No.82001137). We thank Dr. Yulong Li (Peking University) for providing AAV.hSyn.GRAB.NE2h. We thank Mayo Clinic Microscopy and Cell Analysis Core facility for experimental and technical support and members of the Wu Lab for insightful discussions.

## Author contributions

K.H., Y.Y., H.D. and L.-J.W. designed the studies. K.H., A.D.U. and L.-J.W. wrote the manuscript. K.H., T.C. and Y.Y. performed animal surgery, image collection and data analyses. K.H. performed immunofluorescence staining experiments and electron microscope experiments. K.H. and T.X. performed 3D image reconstruction. K.H., Y.Y. and M.H.Y. performed behavioral test and analysis. K.H., Y.Y., Y.L., V.K. and G.W. analyzed EEG data. Funding was obtained and the project supervised by L.-J.W.

## Competing interests

The authors declare no competing interests.

## Supplementary Materials

### Materials and Methods

#### Animals

Male and female mice, 2 to 5 months of age, were used in accordance with institutional guidelines. All experimental procedures were approved by the Mayo Clinic’s Institutional Animal Care and Use Committee (IACUC, protocol #5285-20), and were conducted in accordance with the NIH Guide for the Care and Use of Laboratory Animals. Neuron calcium activity was studied in C57BL/6 wild-type (WT) mice (JAX:000664) using adeno-associated viruses (AAV). Cx3cr1^GFP/+^ (JAX:005582) mice were used to visualize microglia and neuron interactions for in vivo two-photon imaging, where neurons were visualized using AAV transfection. Cx3cr1^GFP/+^ mice were also crossed with PV-Cre knock-in mice (JAX:017320) for microglia and PV neuron interaction studies. Cx3cr1^CreER/+^ (JAX: 021160) mice were bred to Adrb2^fl/fl^ mice (generously provided by Dr. Gerard Karsenty at Columbia University) to generate mice which had β2-adrenergic receptors selectively knocked out of microglia. To activate CreER recombination, tamoxifen (#T5648, MilliporeSigma, Burlington, MA) was injected intraperitoneally 6 weeks before imaging or tissue collection (150 mg/kg, dissolved at 20 mg/ml in corn oil, 5 total injections spaced 48hours apart). To pharmacologically ablate microglia, mice were fed a chow containing PLX3397, a colony-stimulating factor 1 (Csf1) receptor inhibitor, for 3 weeks (600 mg/kg, #C-1271, Chemgood, Henrico, VA).

#### Virus injection and chronic window implantation

Under isoflurane anesthesia (3% induction, 1.5% maintenance,), AAVs were injected into the somatosensory cortex (AP: −4.5, ML: +2.0) using a glass pipette and micropump (World Precision Instruments, Sarasota, FL). AAVs were targeted to layer II/III neurons (DV: −0.2). A 250 nl volume was dispensed at a 40 nl/min rate followed by a 10 min waiting period for diffusion. To image neuronal calcium activity, 250 nL of AAV9.CaMKII.GCaMP6s was injected into the somatosensory cortex (2 mm posterior and 1.5 mm lateral to bregma). To study the relationship between microglia and neuron interactions and neuron calcium activity, we injected a cocktail of three viruses: AAV1.CaMKII.Cre, AAV9.Syn.Flex.GCaMP6s, and AAV9.FLEX.tdTomato into S1 cortex of Cx3cr1^GFP/+^ mice. To visualize microglia and PV boutons simultaneously, we injected AAV9.FLEX.tdTomato into S1 cortex of Cx3cr1^GFP/+^:PV^Cre/+^mice. To image norepinephrine dynamics, rAAV.hSyn.GRAB.NE2h was injected into the somatosensory cortex for in vivo two-photon imaging (*19*). For the detailed information of all AAV vectors used in the study, please see Supplementary Table 1.

Mice were implanted with a cranial window immediately after virus injection as previously described (*37*). A 5 mm circular coverslip was placed above the craniotomy, while a 3mm donut-shaped coverslip was placed within the craniotomy. Coverslips were sealed in place with light-curing dental cement (Tetric EvoFlow, Ivoclar, Buffalo, NY). The skull, excluding the region with the window, was then covered with iBond Total Etch primer (Heraeus, Germany) and cured with LED light. Finally, a head bar (Neurotar, Helsinki, Finland) was attached to the skull using light-curing dental glue, and all exposed skull surface was also covered by dental glue. Mice were allowed to recover from anesthesia on a heating pad for 10 min before they were returned to their home cage. Ibuprofen in drinking water was provided as an analgesic for 48 hours post-surgery.

#### In vivo two-photon imaging

After recovery from virus injection and cranial window surgery (2–4 weeks), mice were trained to move on an air-lifted platform (NeuroTar, Helsinki, Finland) while head-fixed under a two-photon microscope. Training occurred for 30 min/day for the first 3 days following surgery and once per week thereafter. Across all studies, mice were allowed 10 min to acclimate after being placed in the head restraint before imaging began. Mice were imaged using a two-photon imaging system (Scientifica, UK) equipped with a tunable Ti: Sapphire Mai Tai Deep See laser (Spectra Physics, CA, USA). Laser wavelength was tuned to 920 nm to image GCaMP6s fluorescence and Cx3cr1^GFP/+^ microglia, or 950 nm to image tdTomato-labelled neurons and GFP microglia simultaneously. Imaging utilized a 40× water-immersion lens and a 180×180 μm field of view (512×512 or 1024×1024 pixel resolution). The microscope was equipped with a 565 nm dichroic mirror and the following emission filters: 525/50 nm (green channel) and 620/60 nm (red channel) for GFP/tdTomato imaging. The laser power was maintained at 30–40 mW. Imaging in cortex was conducted 50–150 μm beneath the pial surface (Layer I) or 150–200 μm depth (Layer II/III).

To image neuronal calcium activity, WT mice were first imaged under awake baseline conditions (15 min; 150–200 µm depth). A nose cone was then secured against the head-restraint system and the mouse was induced with isoflurane on the platform (3%, up to 60 s). Isoflurane was then maintained at 1.5% for the duration of anesthesia imaging, acquired as two 15 min blocks. Under isoflurane, body temperature was maintained at 37°C using a closed-loop heating system (Physitemp, Clifton, NJ). To end anesthesia, the nose cone was removed, and mice were imaged during this recovery or anesthesia emergence period over four 15 min blocks (60 min emergence). Mouse locomotion was recorded during in vivo calcium imaging using the Mobile HomeCage magnetic tracking system (Neurotar, Helsinki, Finland). To image microglial dynamics, we acquired Z-stacks (8 sections, 2 μm step size) once every minute. To image microglia-neuron interactions, we also acquired Z-stacks (18 sections, 2 μm step size, 512×512-pixel resolution) once every 2-3 minutes.

#### Imaging data analysis

Z-stack images and time-series images were corrected for focal plane displacement using ImageJ (National Institutes of Health, Bethesda, MD, https://imagej.nih.gov/ij/) with the plug-ins TurboReg and StackReg (*38*). To analyze time-series neuronal Ca^2+^ activity, an average intensity image of the entire video was generated for region of interest (ROI) selection. ROIs were manually drawn with the oval selection tool for all neuronal cell somas detected in layer II/III. Neurons that did not show a bright ring and dark nuclei during the awake period (baseline) were excluded from the analysis as they did not express GCaMP6 appropriately. Using these ROIs, mean fluorescent intensity (MFI) values were obtained and converted into ΔF/F. The baseline fluorescence was considered as the lower 25^th^ percentile value across all frames. To classify changes in neuronal activity before and after anesthesia, the following criteria were used: increased (the signal area is more than 1.5 times the baseline), decreased (less than half the baseline). Microglial territories were determined from Z-stack maximum projection images and were calculated as the outermost area of microglia when the distal process tips are connected by a polygon. Microglial contacts were defined as instances where Cx3cr1-GFP signals merge with tdTomato or GCaMP6 signals defining neuronal cell somas in the same focal plane. In classifying neurons by microglial contacts, those with two or more contacts at the end of anesthesia were classified as high contact and those with no or one contact were classified as low contact. A microglial bulbous ending was defined as having a head diameter at the process tip that is at least twice the diameter of the process neck. To determine microglial process motility, maximum z-projection images were used to track primary process tips using the manual tracking plugin in ImageJ. Directness of microglial processes were calculated as Euclidean distance of process tips divided by accumulated distance.

To examine norepinephrine dynamics, MFI of time-series images were measured for the entire field of view and were converted into ΔF/F. The baseline fluorescence was considered as the lower 25^th^ percentile value across all frames.

#### Behavioral tests

To assess mechanical hypersensitivity, von Frey filaments (0.07–2.0 g, North Coast Medical, USA) were applied to the plantar surfaces of the hindpaws before and after isoflurane induction and the 50% withdrawal threshold value was determined using the up–down method (*39*).

#### Immunohistochemistry

Mice were deeply anesthetized with isoflurane then transcardially perfused first with phosphate-buffered saline (PBS), followed by 4% paraformaldehyde (PFA). Brains were post-fixed overnight in 4% PFA and immersed in 30% sucrose for 2 days for cryoprotection. The brains were sectioned into 30 μm slices. After PBS wash, slices were incubated for 1 hour in 5% bovine serum albumin (BSA) and 0.5% Triton X-100 dissolved in PBS (PBST), before being incubated at 4 °C overnight with primary antibodies (anti-IBA1, 1:500, #ab5076 Abcam; anti-NeuN, 1:500, #ab104225 Abcam; anti-VGAT, 1:500, #131 004 Synaptic Systems; anti-VGLUT1, 1:500, #135 304 Synaptic Systems) diluted in PBST. The slices were subsequently incubated with secondary antibodies (anti-goat Alexa 488, anti-rabbit Alexa 647, 1:500, Thermo Fisher Scientific; anti-guinea pig Cy3, 1:500, Jackson ImmunoResearch Labs) at room temperature (RT) for 3 hours. Slices were then mounted on glass slides with DAPI Fluoromount-G (0100-20; Southern Biotech). Fixed tissue was imaged using 63× oil-immersion objective lens with a Zeiss LSM980 Airyscan 2 for confocal imaging or Zeiss ELYRA PS.1 for super-resolution imaging (Carl Zeiss, Oberkochen, Germany). For the detailed information of antibodies used in the study, please see Supplementary Table 2.

#### Quantitative analysis of immunohistochemistry data

In examining the number of bulbous endings on microglia contacting neuronal somata, single plane confocal images were used to count contacts with NeuN-positive neuronal somata. The interaction between bulbous endings and VGAT puncta were characterized from confocal 3D image data (2048×2048 pixels, 0.082 μm/pixel, 0.5 μm Z-step, 20 slices). Volume rendering and visualization were performed using Imaris 9.2 (Oxford Instruments, Abingdon, UK). After applying a threshold (MaxEntropy method) for each channel, bulbous ending structures were classified into four types on interaction: (1) No contact, no merger of IBA1 and VGAT puncta, (2) Contact, under 50% merger of IBA1 and VGAT puncta area, (3) Enwrapped, more than 50% merger of IBA1 and VGAT puncta area, and (4) Engulfment, complete internalization of the VGAT puncta area by IBA1. Co-localization of microglial bulbous-ending and synaptic marker fluorescent signals was used to determine interactions, defined by IBA1 positive area overlapping with VGAT or VGLUT1 positive area. For this analysis, 50×50 pixels around the bulbous ending were cropped from the 63X confocal image, and thresholds were applied using the MaxEntropy or Otsu method for each channel. The co-localized area was calculated in each frame. Microglia density was quantified for each instance in which an IBA1 cell had a DAPI positive nucleus identifiable in the Z stack.

#### Near-infrared branding (NIRB)

We used the NIRB technique to create laser-defined fiducial marks for later electron microscopy identification of cells observed under two-photon microscopy (*40*). After two-photon imaging, mice were perfused with 4% PFA in 0.1 M PBS (pH 7.2). The brain tissue was cut into 1mm thick horizontal sections and post-fixed for 2 h. After fixation, sections were mounted in PBS on a glass slide.

We made NIRB marks on fixed tissue using the two-photon microscope laser. Initially, XYZ images were taken to re-identify the same locations as in vivo imaging. After identification, the laser was tuned to 800nm wavelength (laser power: ∼400 mW in the focal plane) and was focused at 99× through the objective. We made linear NIRB markings along a square path of 150 μm per side surrounding the target microglial cell. The time of laser irradiation was adjusted to ensure that the laser burn did not extend into the target microglia. After NIRB marks were made, sections were post-fixed by immersion in 2% glutaraldehyde + 2% paraformaldehyde in 0.1 M cacodylate buffer containing 2 mM calcium chloride at 4 °C for at least 24 hours.

#### Serial block-face scanning electron microscopy

Samples for serial block-face scanning electron microscopy (SBF-SEM) were prepared as reported (*40*). Briefly, NIRB marked samples were rinsed in 0.1 M cacodylate buffer and placed in 2% osmium tetroxide + 1.5% potassium ferracyanide in 0.1 M cacodylate, then washed with double distilled water (ddH_2_O) incubated at 50°C in 1% thiocarbohydrazide, incubated again in 2% osmium tetroxide in ddH_2_O, rinsed in ddH_2_O and then placed in 2% uranyl acetate overnight. The next day sample were rinsed in ddH_2_O, incubated with Walton’s lead aspartate, dehydrated through an ethanol series, and embedded in Embed 812 resin (Electron Microscopy Sciences, Hatfield, PA). Based on ferrocyanide-reduced osmium-thiocarbohydrazide-ferrocyanide-reduced osmium (rOTO) stains (*41*), this procedure introduces a considerable amount of electron-dense, heavy metals into the sample to provide the contrast necessary for SBF-SEM.

To prepare the embedded sample for placement into the SBF-SEM, a ∼1.0 mm^3^ piece was trimmed of any excess resin and mounted on an 8 mm aluminum stub using silver epoxy Epo-Tek (EMS, Hatfield, PA). The mounted sample was then carefully trimmed into a smaller ∼0.5 mm^2^ column using a Diatome diamond trimming tool (EMS, Hatfield, PA) and vacuum sputter coated with gold palladium to help dissipate electrostatic charge. Sectioning and imaging of the sample was performed using a VolumeScope 2 SEM^TM^ (Thermo Fisher Scientific, Waltham, MA). Imaging was performed under high vacuum/ low water conditions with a starting energy of 1.8 keV and beam current of 0.10 nA. Sections of 50 nm thickness were cut allowing for imaging at 8 nm x 8 nm x 50 nm spatial resolution.

#### 3D image segmentation

Image analysis, including registration, volume rendering, and visualization were performed using *ImageJ*, *Amira* (Thermo Fisher Scientific), and *Reconstruct* (*42*) software packages. Cell morphology and ultrastructure were identified and segmented in every 50 nm thick section using two-photon images as a reference. In segmenting microglial morphology, cell soma were first identified using established criteria, which include relatively dark nuclei with clumped chromatin, narrower space between nuclear membranes, a dark cytoplasm and processes containing elongated endoplasmic reticulum (*43*). The processes of these microglia were then tracked in across 3D images to identify bulbous endings associated with specific elements of the neuron.

Synapses were characterized by a presynaptic element or varicosity that contains synaptic vesicles and forms one or more electron-dense junctions with an axon terminal or cell body of the postsynaptic neuron (*44*). Synapses in displacement were defined as one in which the bouton is adjacent to the neuronal cell body with microglial processes between them (*45, 46*). If a structure other than microglia was present in the gap, it was excluded because it was determined that they had not originally formed synapses there. The PV-bouton structure was estimated based on fluorescence images of tdTomato, which virally labeled PV-boutons in two-photon microscopy. This ultrastructure was determined by segmentation of the actual electron microscope images.

#### EEG recording and analysis

Under Isoflurane anesthesia (3% induction, 1.5% maintenance), a cortical electrode (Plastics One, Roanoke, VA) was inserted into layer II/III of the parietal cortex (200 μm depth). Electrodes were secured by super glue (Loctite 454, Westlake, OH) against three anchor screws. Animals were used for EEG recordings after 1–2 weeks of recovery. During recordings, electrodes were connected via flexible tether to a rotating commutator (Plastics One, Roanoke, VA) then connected to an EEG100C amplifier (BioPac Systems, Goleta, CA), which filtered signals below 1 Hz and above 100 Hz.

To consistent with two photon imaging strategy, mice were recorded 30 minutes of baseline EEG under head-restraint system, then mouse was induced with isoflurane on the platform (3% induction, 1.5% maintenance). Post-processing of the intracranial EEG (iEEG) data was performed using custom algorithms (Matlab version 2019b, MathWorks, Natick, MA) for characterizing the changes of iEEG power in spectral bands of interest (Delta (1–3 Hz), Theta (3–9 Hz), Alpha (8– 12 Hz), Beta (15–30 Hz),) as described previously (*47*). Absolute and relative power spectral densities (PSD) across frequency bands were calculated using 10 second windows (epochs) and the average of PSD from all epochs was calculated as a representative value of PSD in each phase of experiment for each subject. Absolute PSD refers to the absolute power (V_rms2_/Hz), where V_rms_ is the root-mean-square voltage of the subject’s iEEG, while relative PSD refers to the percentage of power a frequency band (such as beta) compared to the sum of all frequency bands.

## Data analysis and statistics

Data were analyzed using GraphPad Prism 8 software (GraphPad Software Inc., La Jolla, CA). All data are presented as the mean ± SEM. Statistical significance was determined using either unpaired or paired t-tests, one-ANOVA followed by Tukey’s or Sidak’s post-hoc test as indicated in either the results or figure legends.

---

**Figure S1:**
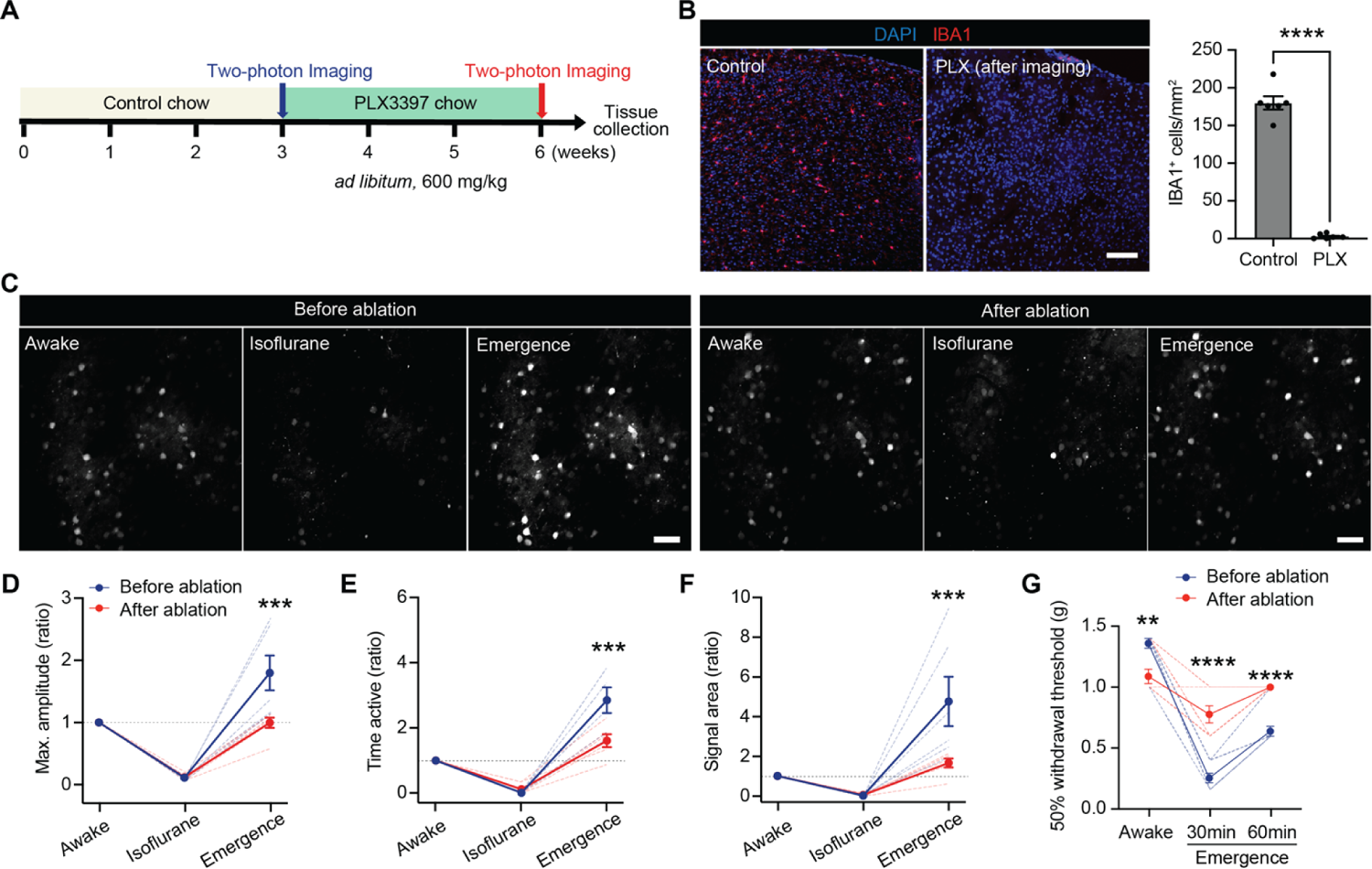
Microglial depletion abolishes the increase of neuronal activity during emergence. (A) Experimental timeline of microglia ablation and two-photon imaging of neuronal Ca^2+^ activity before and after general anesthesia. (B) IBA1 immunostaining of microglial density in mice fed with a control chow (Intact, left panel) and a PLX3397-containing chow (PLX, middle panel) in the cortex (blue for DAPI and red for IBA1). Graph shows significantly decreased IBA1^+^ cells in PLX-treated mice (right). Scale bar: 50 μm. Unpaired t-test, ****p < 0.0001. (C) Representative images of neuronal Ca^2+^ activity across experimental phases in mice before (left panels) and after microglia ablation (right panels). Scale bar: 50 μm. (D) Graph shows maximum amplitude (ratio) of neuronal Ca^2+^ active time before and after ablation. n = 6 mice each group. (E) Graph shows time active (ratio) of neuronal Ca^2+^ activity before and after ablation. (F) Graph shows ΔF/F signal area (ratio) of neuronal Ca^2+^ activity before and after ablation. (G) Graph shows reduced mechanical hypersensitivity during the emergence from anesthesia (30 min) in microglia ablated mice. n = 10 before ablation, n = 9 after ablation mice. In all graphs, each point or dashed lines indicates data from an individual animal (B, D, E, F and G), while columns or solid lines and error bars show the mean ± SEM. Two-way ANOVA followed by Tukey post-hoc test; **p < 0.01, ***p < 0.001 and ****p < 0.0001.

**Figure S2:**
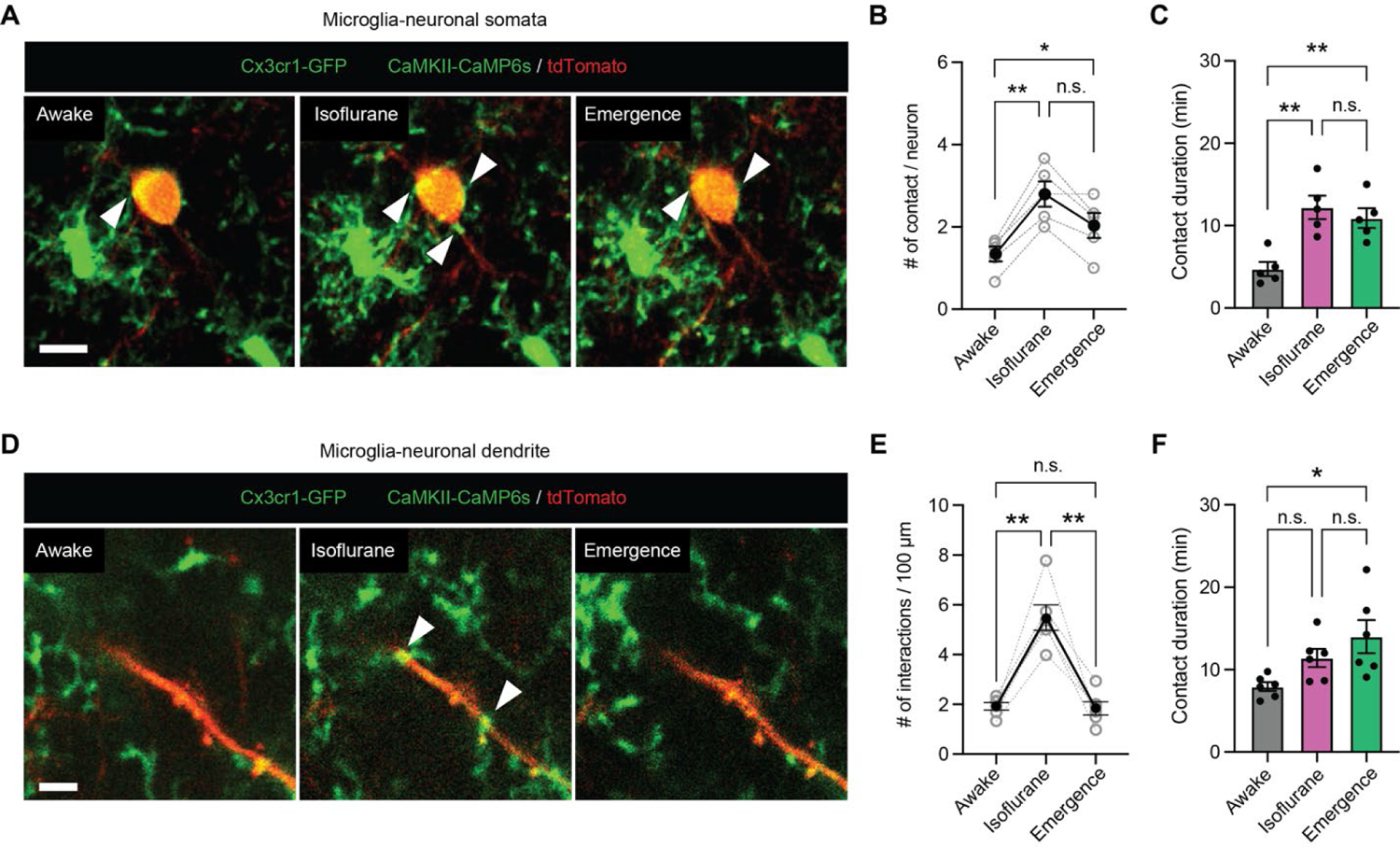
Increased contacts between microglia and neuronal somata or dendrites in response to general anesthesia. (**A**) Two-photon imaging of microglia (Cx3cr1^GFP/+^, green) and neurons co-labelled with GCaMP6s (green) and tdTomato (red). Representative time series images of interaction between microglial bulbous endings (green) and neuronal soma (red) during anesthesia. White arrowheads indicate contact sites. Scale bar: 5 μm. (**B**) Graph shows that number of microglia-neuronal soma contact sites was significantly increased during anesthesia. N = 5 imaging fields from 5 mice. (**C**) Duration of microglial interaction before (Awake), during (Isoflurane) and after anesthesia (Emergence). N = 5 mice. (**D**) Representative time series images of interaction between microglial bulbous endings (green) and neuronal dendrites (red) before, during, and after anesthesia. Microglia-neuronal dendrite interaction was increased during anesthesia (indicated by white arrowheads). Scale bar: 5 μm. (**E**) Graph shows number of microglial interactions per 100 μm length of dendrite. N = 6 imaging fields from 3 mice. (**F**) Graph shows duration of observed microglia-dendrite interactions before, during, and after anesthesia. N = 6 imaging fields from 3 mice. Dashed lines and dots indicate data from an individual animal, solid lines, columns and error bars show the mean ± SEM (**B**, **C**, **E** and **F**). One-way ANOVA followed by Tukey’s post-hoc test; n.s., not significant; *p < 0.05 and **p < 0.01.

**Figure S3:**
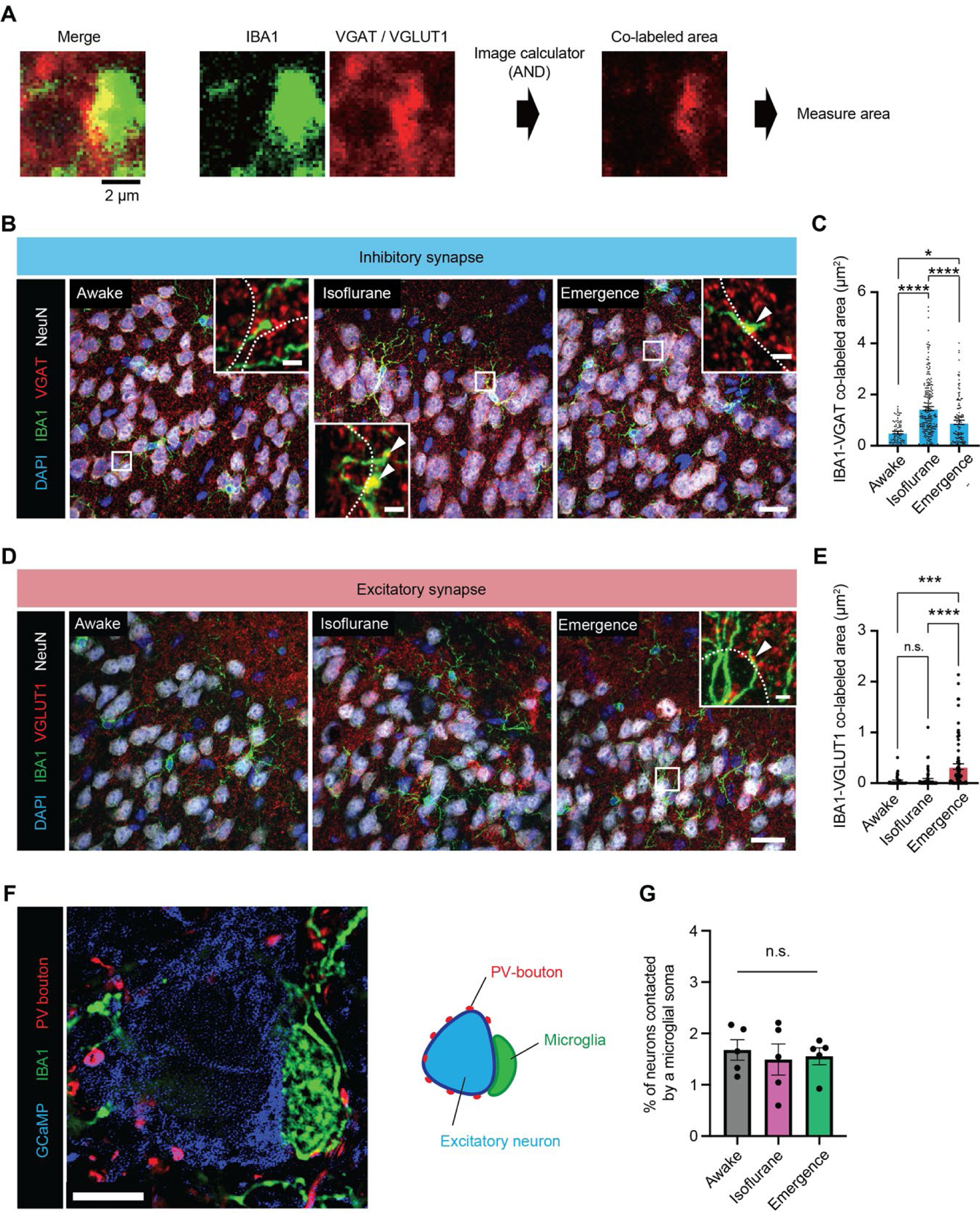
Increased colocalizations between microglial bulbous endings with VGAT puncta during anesthesia and emergence. (**A**) Schematic diagrams for colocalization analysis of microglial bulbous ending (IBA1, green) and a synapse marker (red, VGAT or VGLUT1) in confocal microscopy images. (**B**) Representative images of microglia (IBA1, green) and GABAergic inhibitory synapses (VGAT, red) before, during, and after anesthesia. (**C**) Quantification of IBA1^+^ and VGAT^+^ colocalized area (μm^2^) in imaging fields at each time point. n = 80 (Awake), n = 215 (Isoflurane), n = 124 (Emergence) bulbous endings. (**D**) Representative images of microglia (IBA1, green) and glutamatergic excitatory synapses (VGLUT1, red) before, during, and after anesthesia. (**E**) Graph shows IBA1^+^ and VGLUT1^+^ colocalized area (μm^2^) in imaging fields at each time point. n = 37 (Awake), n = 58 (Isoflurane), n = 80 (Emergence) bulbous endings. (**F**) Confocal imaging of representative soma-soma interaction between GCaMP^+^ neurons (blue) and microglia (green). PV^+^ boutons (red) were also shown. Diagram illustrates the observed phenomenon (right). (**G**) Graph shows percentage of soma-soma interaction between microglia and neurons for all neuronal populations in the imaging field before, during, and after anesthesia. n = 5 mice. Each point indicates data from an individual bulbous ending interaction (**C and E**) or mouse (**G**) while bars and error bars indicate the mean ± SEM. One-way ANOVA followed by Tukey’s post-hoc test; n.s., not significant; *p < 0.05, **p < 0.01 and ***p < 0.001. Scale bar: 20 μm or 2 μm (inset) (**B** and **D**), 5 μm (**F**).

**Figure S4:**
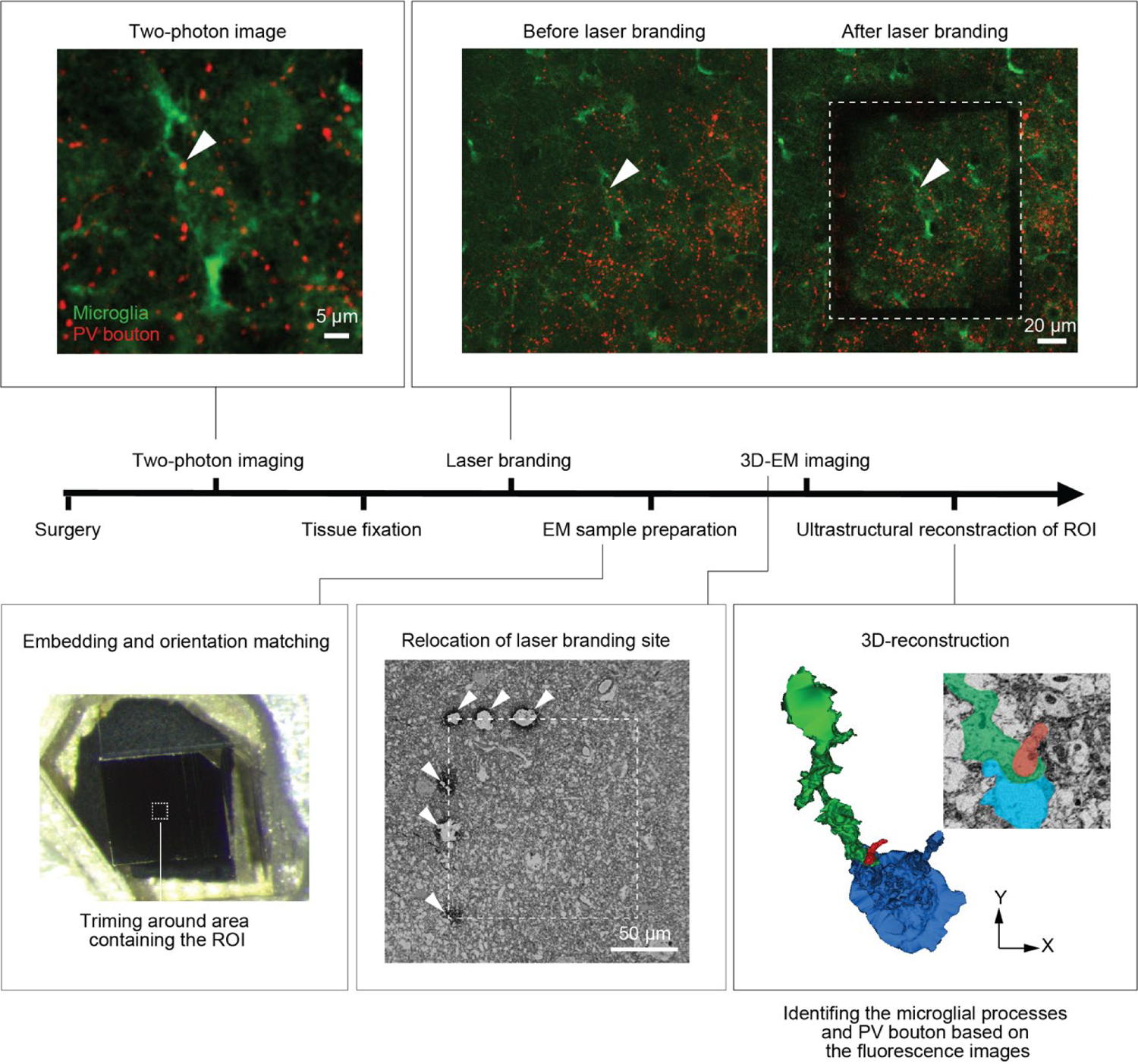
Near-infrared branding to correlate two photon images with electron microscopy and study microglia-neuron interaction Timeline of experimental procedures necessary to capture an EM volume from the site of two-photon live imaging. Following *in vivo* two-photon imaging, brain tissue was fixed. We then re-identified the same region of imaging in fixed tissue under the two-photon microscope and performed near-infrared branding to create defined fiducial marks. This region was further cut down and processed for EM imaging, including tissue embedding and dehydration. Serial block-face scanning electron microscopy (SBF-SEM) was performed on sequential sections using the fiducial marks from laser branding to ensure proper tissue orientation. The image was reconstructed for microglial bulbous ending interactions with PV boutons and neuronal somata at the ultrastructural level. For details, please see the methods section.

**Figure S5:**
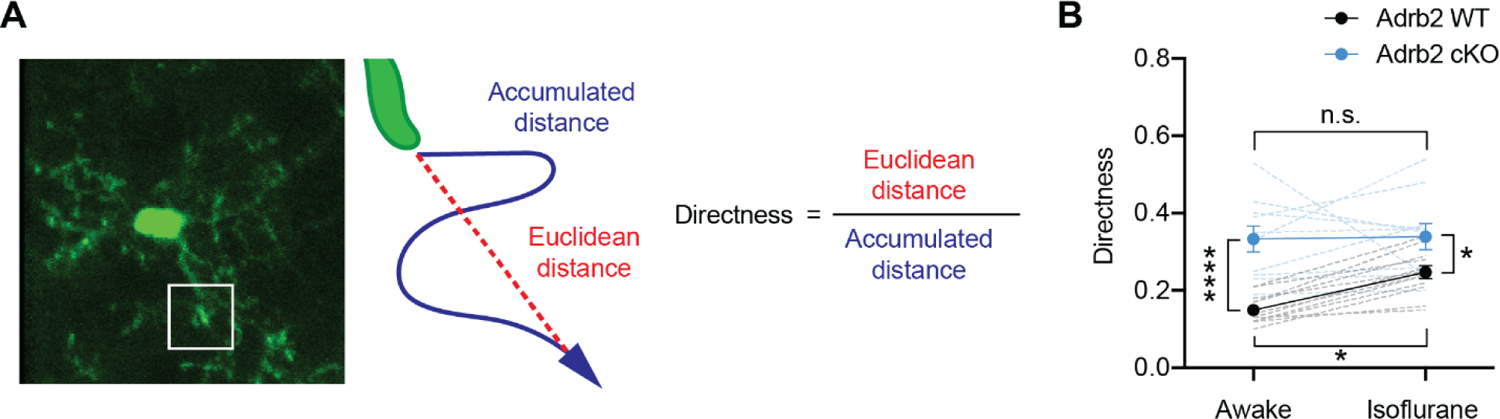
The increased directness of microglial process movement in response to anesthesia. (**A**) A representative image of microglia with dynamic processes (left). The illustration describes the calculation of microglial process moving outward in its trajectory. Microglial process directness was calculated as the Euclidean distance of the process tips divided by its accumulated distance. (**B**) Graph shows increasing process trajectory directness during anesthesia in Adrb2 WT mice but not in Adrb2 cKO mice. Each dashed lines indicates data from individual microglia: n = 12 WT microglia and n = 10 cKO microglia from 5 mice. Two-way ANOVA followed by Sidak’s post-hoc test; n.s., not significant; *p < 0.05 and ****p < 0.0001.

**Figure S6:**
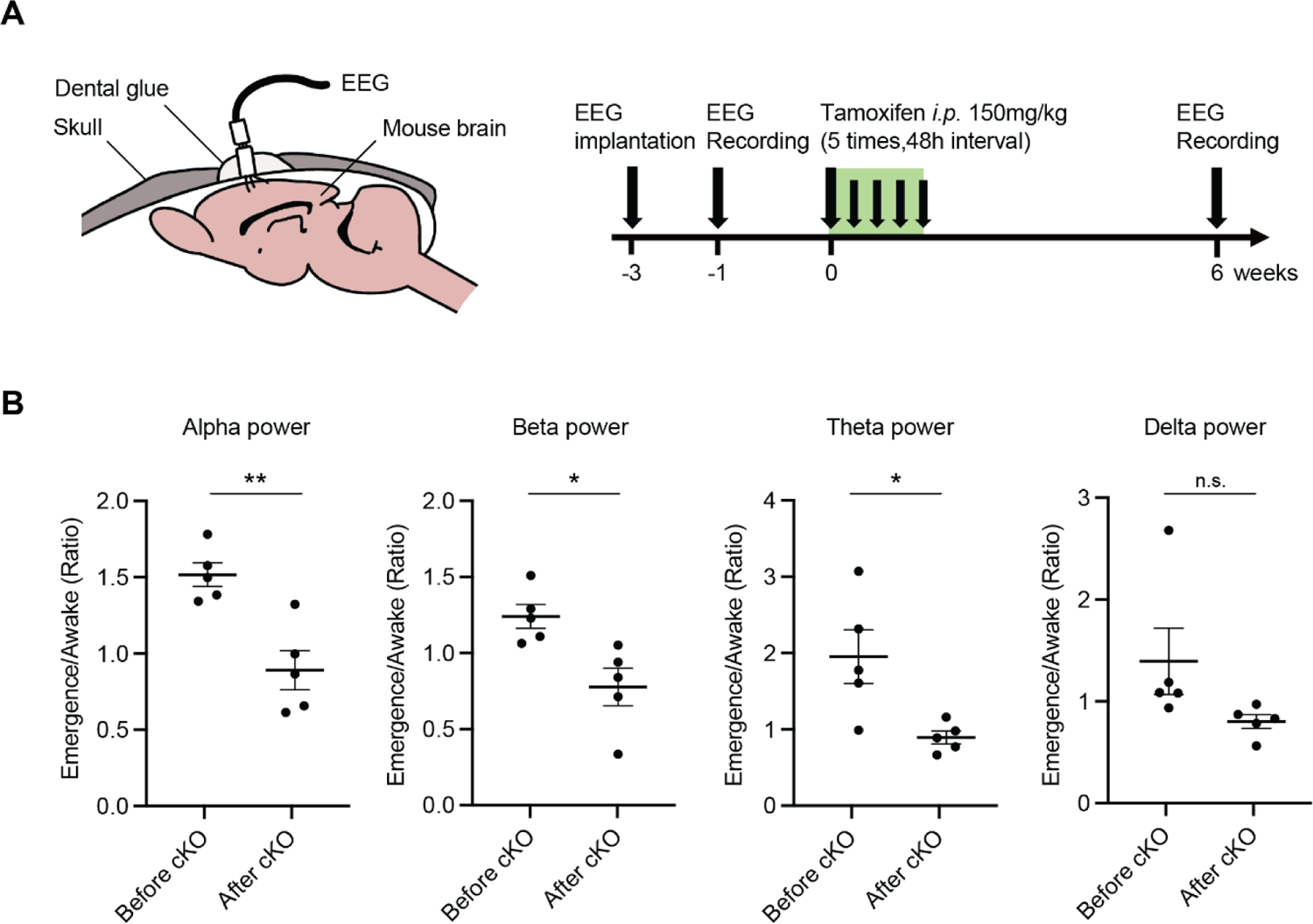
Adrb2 deletion shifts in EEG power during emergence (**A**) Diagrams of EEG electrode implantation and the study timeline. (**B**) Power in the alpha, beta, theta, and delta frequency bands is expressed as a ratio of power in the emergence phase relative to the awake phase before and after recombination for Adrb2 cKO. n = 5 mice. Paired t-test; n.s., not significant; *p < 0.05, **p < 0.01.

**Table S1.** List of AAV vectors used in the study.

| **Name** | **source** | **catalog no.** | **RRID** |
| --- | --- | --- | --- |
| AAV.CamKII.GCaMP6s.WPRE.SV40 | Addgene | 107790-AAV9 | Addgene 107790 |
| AAV.CamKII 0.4.Cre.SV40 | Addgene | 105558-AAV1 | Addgene 105558 |
| pAAV.Syn.Flex.GCaMP6s.WPRE.SV40 | Addgene | 100845-AAV9 | Addgene 100845 |
| pAAV-FLEX-tdTomato | Addgene | 28306-AAV9 | Addgene 28306 |
| AAV9-hsyn-NE2h | WZbioscience | YL003011-AV9 | n.a. |

**Table S2.**
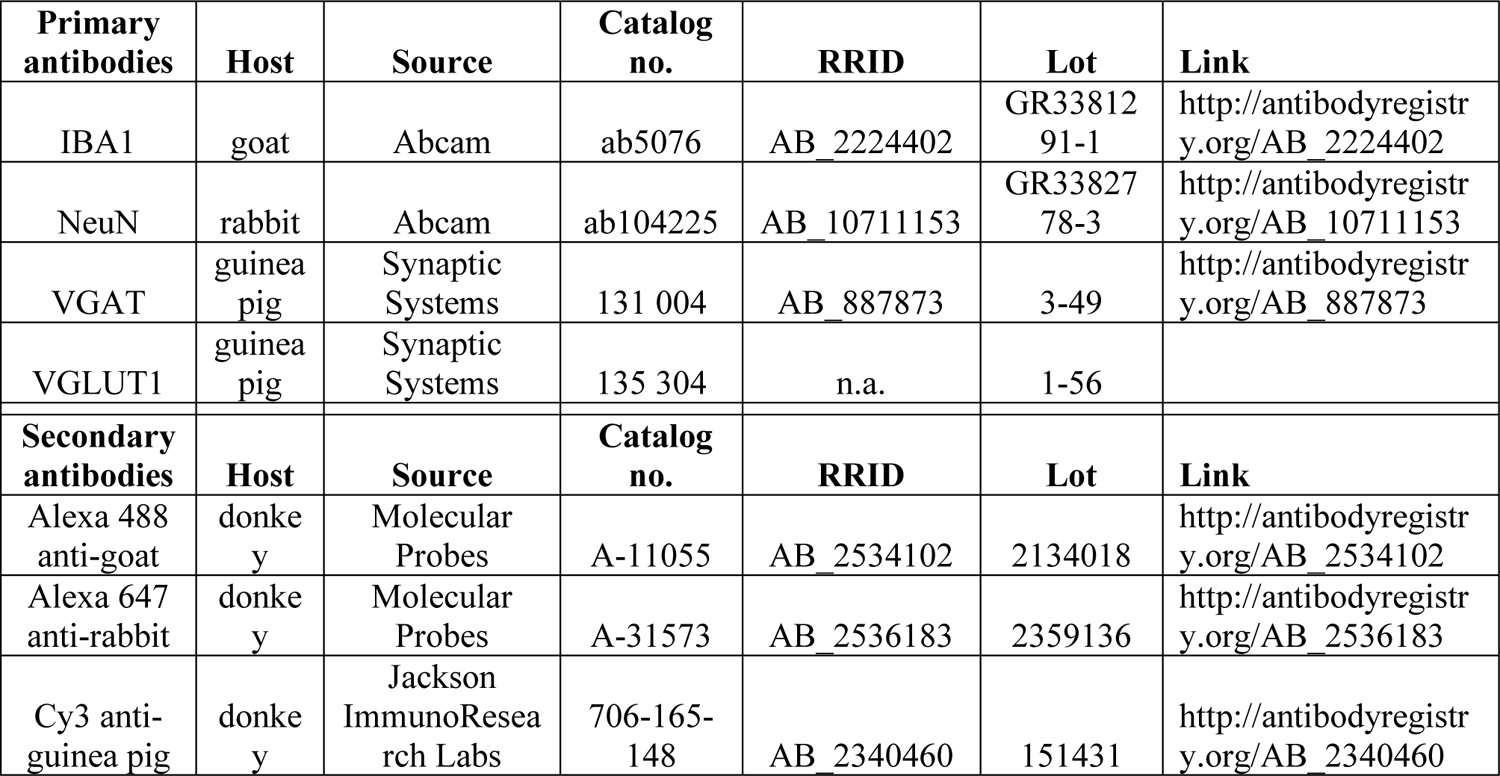
List of antibodies used in the study.

| **Primary antibodies** | **Host** | **Source** | **Catalog no.** | **RRID** | **Lot** | **Link** |
| --- | --- | --- | --- | --- | --- | --- |
| IBA1 | goat | Abcam | ab5076 | AB_2224402 | GR33812<br>91-1 | http://antibodyregistry.org/AB_2224402 |
| NeuN | rabbit | Abcam | ab104225 | AB_10711153 | GR33827<br>78-3 | http://antibodyregistry.org/AB_10711153 |
| VGAT | guinea pig | Synaptic Systems | 131 004 | AB_887873 | 3-49 | http://antibodyregistry.org/AB_887873 |
| VGLUT1 | guinea pig | Synaptic Systems | 135 304 | n.a. | 1-56 |  |
| **Secondary antibodies** | **Host** | **Source** | **Catalog no.** | **RRID** | **Lot** | **Link** |
| Alexa 488 anti-goat | donkey | Molecular Probes | A-11055 | AB_2534102 | 2134018 | http://antibodyregistry.org/AB_2534102 |
| Alexa 647 anti-rabbit | donkey | Molecular Probes | A-31573 | AB_2536183 | 2359136 | http://antibodyregistry.org/AB_2536183 |
| Cy3 anti-guinea pig | donkey | Jackson ImmunoResearch Labs | 706-165-148 | AB_2340460 | 151431 | http://antibodyregistry.org/AB_2340460 |

## Supplementary movies

**Movie S1.** Related to Fig. 1C. *In vivo* Ca^2+^ imaging of layer II/III excitatory neurons in somatosensory cortex before, during, and after isoflurane anesthesia. Calcium activity in excitatory neuronal somata increases after the cessation of anesthesia compared to the awake period.

**Movie S2.** Related to Fig. 1L. *In vivo* two-photon imaging of microglial dynamics in Cx3cr1^GFP/+^ mice before, during, and after isoflurane anesthesia. A raw image, single cell, and territory area are displayed after thresholding, in that order. Microglial territory area and number of bulbous endings increase during anesthesia.

**Movie S3.** Related to fig. S1C. *In vivo* Ca^2+^ imaging of layer II/III excitatory neurons in somatosensory cortex before and after microglia ablation. Emergence-induced hyperactivity was suppressed after microglia ablation.

**Movie S4.** Related to Fig. 2A. *In vivo* two-photon imaging of microglia-neuronal somata interactions. (Microglia: GFP, green; CaMKII^+^ neuron: tdTomato, red, and GCaMP6s, green). Arrowheads indicate bulbous endings in contact with the neuronal somata.

**Movie S5.** Related to Fig. 3H. *In vivo* two-photon time-lapse imaging of microglia-PV bouton interactions. (Microglia: GFP, green; CaMKII^+^ neuron: GCaMP6s, green; PV bouton: tdTomato, red). White circle outlines neuronal somata. Arrowheads indicate contact sites. PV bouton remains after retraction of the microglial process.

**Movie S6.** Related to Fig. 4D. The 3-dimensional serial reconstruction of SBF-SEM images shows that the process tips of microglia insert themselves into a space between neuronal somata and PV boutons. The first half of the video displays microglia (green), PV boutons (red) and neuronal somata (blue), while the second half only shows microglia and PV boutons for simplicity.

**Movie S7.** Related to Fig. 5B. *In vivo* two-photon imaging of NE release in WT mice expressing a genetically encoded NE sensor on neuronal membranes. NE concentration decreased in the parenchyma of Layer II/III somatosensory cortex during anesthesia but increased above awake baseline levels in the first 30 minutes of emergence.

**Movie S8.** Related to Fig. 5E. *In vivo* two-photon imaging of microglial dynamics in Adrb2 WT and Adrb2 cKO mice before, during and after isoflurane anesthesia. Increased microglial process dynamics were lost in Adrb2 cKO mice.

**Movie S9.** Related to Fig. 5, J and K. *In vivo* Ca^2+^ imaging of layer II/III excitatory neurons under awake baseline, anesthesia, and emergence before and after Adrb2 cKO. Emergence-induced hyperactivity exists prior to knockout of Adrb2 but is lost in Adrb2 cKO mice.

## Notes

### Competing Interest Statement

The authors have declared no competing interest.

